# Tuning ice thickness using the chameleon for high-quality cryoEM data collection

**DOI:** 10.1101/2024.05.01.592094

**Authors:** Kelly L. McGuire, Brian D. Cook, Sarah M. Narehood, Mark A. Herzik

## Abstract

Advances in single-particle cryogenic electron microscopy (cryoEM) now allow for routine structure determination of well-behaved biological specimens to high-resolution. Despite advances in the electron microscope, direct electron detectors, and data processing software, the preparation of high-quality grids with thin layers of vitreous ice containing the specimen of interest in random orientations remains a critical bottleneck for many projects. Although numerous efforts have been dedicated to overcoming hurdles frequently encountered during specimen vitrification using traditional blot-and-plunge specimen preparation techniques, the development of blot-free grid preparation devices provide a unique opportunity to carefully tune ice thickness, particle density, and specimen behavior during the vitrification process for improvements in image quality. Here, we describe critical steps of high-quality grid preparation using a SPT Labtech chameleon, evaluation of grid quality/ice thickness using the chameleon software, high-throughput imaging in the electron microscope, and recommend steps for troubleshooting grid preparation when standard parameters fail to yield suitable specimen.

**Video Link:** Contents of this manuscript are available as a video tutorial. This video can be found **[here](https://www.youtube.com/watch?v=F44ug__WdUU&feature=youtu.be)**

## 1 Introduction

Technological advances in cryogenic electron microscopy (cryoEM) have resulted in a near exponential increase year over year in the number of cryoEM structures being released. Although these technologies have improved the quality of the images being recorded[1–5], the throughput of data collection and data processing[6–10], as well as the quality of the resulting three-dimensional (3-D) reconstructions, these advances do not mitigate the oft-encountered difficulties in EM grid preparation. Indeed, the field of cryoEM still relies heavily on the traditional blot-and-plunge method for vitrifying samples that has remained essentially unchanged since the 1980s.[11–14] This method consists of applying a small volume of sample (e.g., 2-3 *µ*L) to the surface of an EM grid consisting of a film (e.g., copper, gold, etc.) with holes overlaid on a metal support mesh (e.g., copper, gold, etc.). Filter paper is then used to remove excess sample by directly contacting the surface of the EM grid, leaving a thin layer of sample that is subsequently plunge frozen into liquid ethane cooled by liquid nitrogen. This vitrified specimen can then be transferred to the transmission electron microscope (TEM) for image recording.[15]

While some instrumentation has been developed to automate aspects of the vitrification process, many of these devices still require the use of filter paper to remove excess specimen from the grid surface, resulting in inconsistent thickness of the vitrified layer, uneven distribution of ice thickness across the EM grid, and low reproducibility across different grids.[16–19] This unpredictability is further compounded by the deleterious effects from interactions with the air-water interface, resulting in inconsistent specimen behavior in ice.[20,21] As a result of these disadvantages, several methods were developed to eliminate the physical blotting component of EM grid preparation, yielding alternative so-called “self-blotting” approaches.[22–24] These instruments – including the original Spotiton and subsequent chameleon (SPT Labtech) – utilize a piezo-electric inkjet dispensing head to deliver small droplets (e.g., 10s of picoliters) to an EM nanowire grid that self-wicks solution while being plunge frozen.[25–28] This design affords several advantages compared to traditional blotting, including the ability to establish a clear gradient of ice thickness across the grid surface in a so-called “stripe”. This provides the EM practitioner the ability to systematically screen particle distribution and quality across the ice thickness gradient in a high-throughput manner.

In this article, we will describe critical steps of high-quality grid preparation using a SPT Labtech chameleon. This includes initialization of the instrument for grid preparation, proper grid handling techniques to ensure optimal downstream operation, evaluation of the sample dispensing results, and modifying dispenser parameters to obtain the desired ice thickness. We detail methods for tuning the ice thickness based on the sample and discuss high-throughput screening of these grids to evaluate grid quality and particle behavior, methods for identifying areas of thin ice, and how to modify the chameleon parameters during subsequent grid freezing sessions. Additionally, we detail the contribution of ice thickness to particle density and image quality and how to modify the chameleon when standard parameters fail to provide suitable results. Finally, we have developed a comprehensive video protocol that can be used alongside this written protocol for the creation of high quality grids.

## II Protocol

### A Initialize the chameleon and Prepare Grids for Sample Vitrification

This section provides detailed instructions for initializing the chameleon vitrification system and preparing grids for sample vitrification, with an estimated operational time of 15-20 mins. Key steps include filling the wash pump supply, humidifier, and dispenser supply bottles, ensuring appropriate fluid levels in the various solution bottles (**Figure 1**). It is critical to follow specific guidelines for filling these bottles to avoid issues such as clogging and improper dispensing due to contaminants. Additional steps involve setting up the dispenser system, including a clean tweezer vial with ethanol, loading a sample vial properly to prevent air bubbles, and calibrating the system. The section emphasizes cleanliness and proper handling to maintain the system’s integrity and functionality.

**Figure 1:**
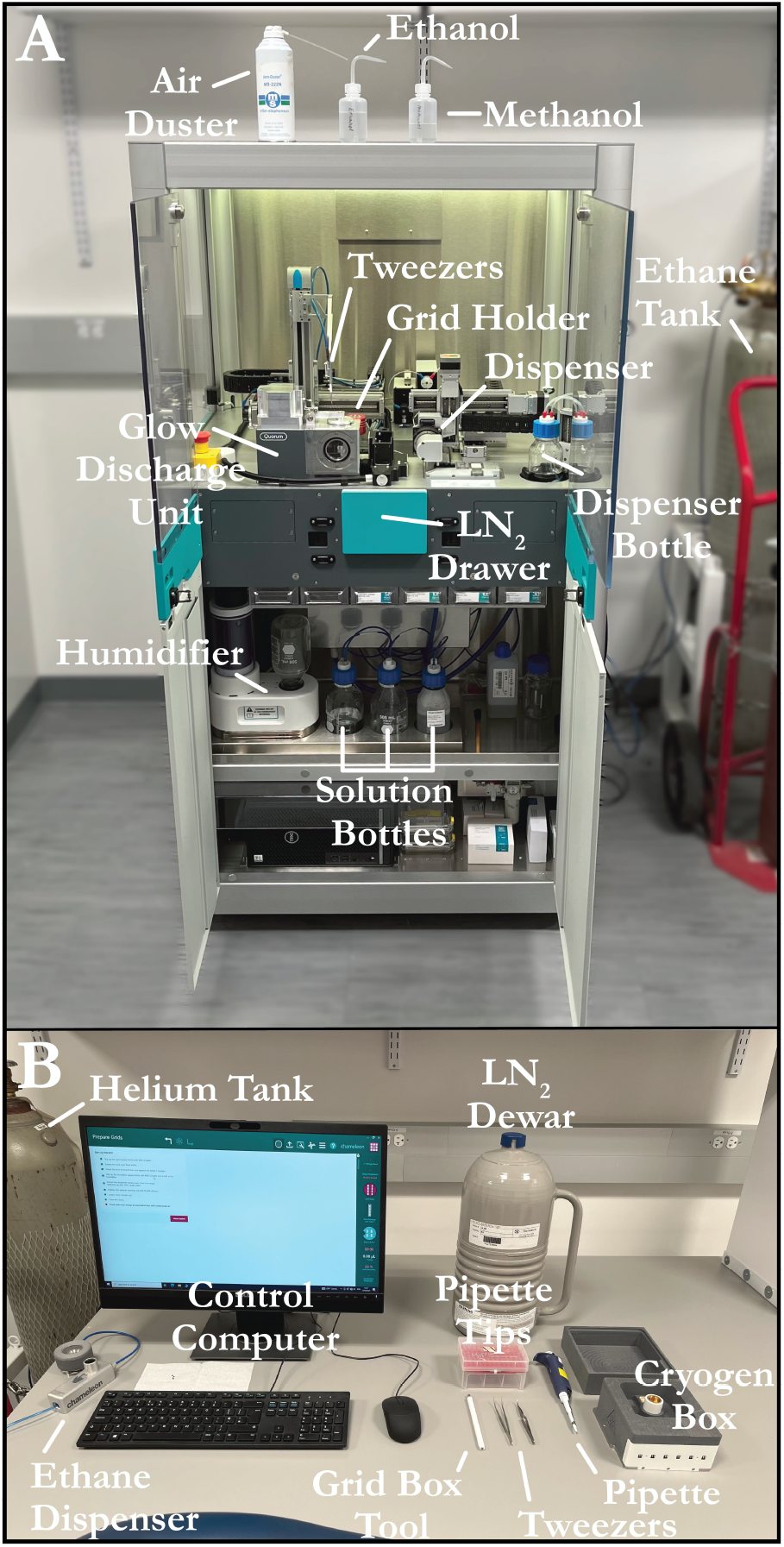
Chameleon layout and required materials. A) A staged chameleon with all doors open and the necessary components. B) The staged chameleon computer workstation. Labels depict the necessary equipment for operating the chameleon.

A.1 On the PC attached to the chameleon, open the chameleon software and verify that the system monitoring lights turn green. Once the chameleon® software has finished starting, a checklist will appear on the <u>Prepare Grids</u> page.
A.2 Fill the wash pump supply bottle with ultrapure water to at least 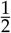 full.
  A.2.1 **NOTE:** Ultrapure water must be used to prevent debris and contaminants from entering the sys-tem that could clog the dispenser and/or alter dispensing properties.
A.3 Ensure the used wash fluid bottle is empty and the tubing is at the bottom of the bottle.
  A.3.1 **NOTE:** The wash fluid bottle should be emptied by the user at the end of each chameleon session (detailed further in **Section D.20**). If this has not been performed, empty the bottle. If needed, adjust the tubing prior to starting the session.
A.4 Check that the level of the ChamClean® bottle is at least 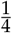 full and the tubing is at the bottom of the bottle.
  A.4.1 **NOTE:** Each session uses approximately 5% of the ChamClean® solution.
  A.4.2 If needed, adjust the tubing prior to starting the session.
A.5 Ensure the level of the humidifier supply bottle is filled to 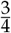 full of ultrapure water. Connect the humidifier bottle to the humidifier and gently rock the bottle until air bubbles stop to make sure it filled the humidifier completely.
  A.5.1 **NOTE:** When filling the bottle, leave an air gap of 50-75 mm between the top of the bottle and the top of the water. If the bottle is filled to the top without leaving an air gap, the water will not transfer to the humidifier due to lack of pressure difference. The air gap also prevents water from flowing from the humidifier back into the bottle.
A.6 Ensure the dispenser supply water level is within the white vertical line indicated on the side of the dispenser supply water bottle and the tubing is at the bottom of the bottle.
  A.6.1 **NOTE:** The bottom and top of the vertical white line indicate the minimum and maximum supply water level range, respectively.
  A.6.2 **NOTE:** Only fill with HPLC-grade water to prevent debris and contaminants from entering the system that could clog the dispenser and/or alter dispensing properties.
A.7 Place a new tweezer cleaning vial to the right of the red grid holder and fill to the top with pure ethanol using the supplied squirt bottle.
  A.7.1 **NOTE:** It is recommended to replace this vial only once per week. During a new session initialization, the current tweezer cleaning vial should be air dusted.
A.8 Load a clean sample vial into the sample holder.
  A.8.1 **NOTE:** It is recommended that the vial be “gently dropped” from about 1-2 mm from above the holder location to allow proper seating of the sample vial in the holder. Do not press the sample vial into the holder. This may set the sample vial too low for proper aspiration, thus introducing air bubbles into the dispenser.
  A.8.2 **NOTE:** In **Section B.9** the user will add a sample to the sample vial. It is recommended to air dust the vial before adding a sample to eliminate possible contaminants.
A.9 The <u>Prepare Grids</u> page includes the “Home System” button. Press this button to calibrate the system by zeroing out the position of the tweezers as well as the liquid nitrogen (LN_2_) drawer.
  A.9.1 **NOTE:** The monitoring lights will turn white and the doors to the chameleon will remain locked until the software has advanced to the <u>Dispenser Setup</u> page.
A.10 The software will advance to the <u>Select Mode</u> page. Enter the operator’s name (or initials) and enter a name for the project.
  A.10.1 **NOTE:** This allows for tracking of dispenser health and functionality across different samples and time.
  A.10.2 **OPTIONAL:** If this session follows a previous user or a previous sample dispensing, it is recommended to perform a thorough cleaning of the dispenser before moving forward. The <u>Select Mode</u> page includes the “Clean Dispenser” button which will perform an additional dispenser cleaning to remove residue from the previous sample.
A.11 Click the “Prepare Grids” button to advance to the <u>Helium Sparge</u> page.
A.12 Open the helium tank valves and click the “Sparge” button to sparge the dispenser bottle. This process allows helium gas to flow through the bottle, degassing the solution. This takes approximately 2 mins. Once finished, close the helium tank valves.
  A.12.1 **NOTE:** The regulator attached to the helium tank should have an outlet pressure reading of 30 psi. Adjust if needed to obtain the proper pressure for ideal operation.
  A.12.2 **CAUTION:** Please consult your institution’s Environmental Health and Safety Guidelines for safe location and operation of a helium tank.
A.13 Click the “Continue” button to advance to the <u>Dispenser Setup</u> page. The system monitoring lights will turn green, and the front main doors will unlock.
A.14 Open the chameleon doors and place a new methanol vial. The methanol vial should be air dusted to eliminate contaminants and filled to the top with methanol using the supplied squirt bottle.
A.15 Click the “Setup” button to initialize the dispenser fluid system. The main doors will lock, and the monitoring lights turn white. Video of the dispenser initialization will show the backflushing process. The user should watch for particulates or debris in the dispenser tip. The “Setup” button will be grayed out during this process. When finished, the main doors will unlock, and the monitoring lights turn green. This process will take approximately 3 mins.
  A.15.1 **NOTE:** If particulates or debris are observed, follow troubleshooting instructions in *Troubleshooting* **Section G.2**.
  A.15.2 **OPTIONAL:** Open chameleon doors and refill the methanol vial to the top with methanol. Close the main doors.
A.16 Click the “Test Dispenser” button. This will advance to the <u>Test Dispenser</u> page and the system will run an automatic initial dispense.
A.17 Click the “Automatic Test” button to run multiple dispenser tests. The software will analyze the results and if successful, the “Dispenser tested ok” message will appear.
  A.17.1 **NOTE:** The default dispenser settings established during the installation should yield four to five separate droplets with the droplet farthest from the dispenser tip landing near the bottom of the image (**Figure 2**).

**Figure 2:**
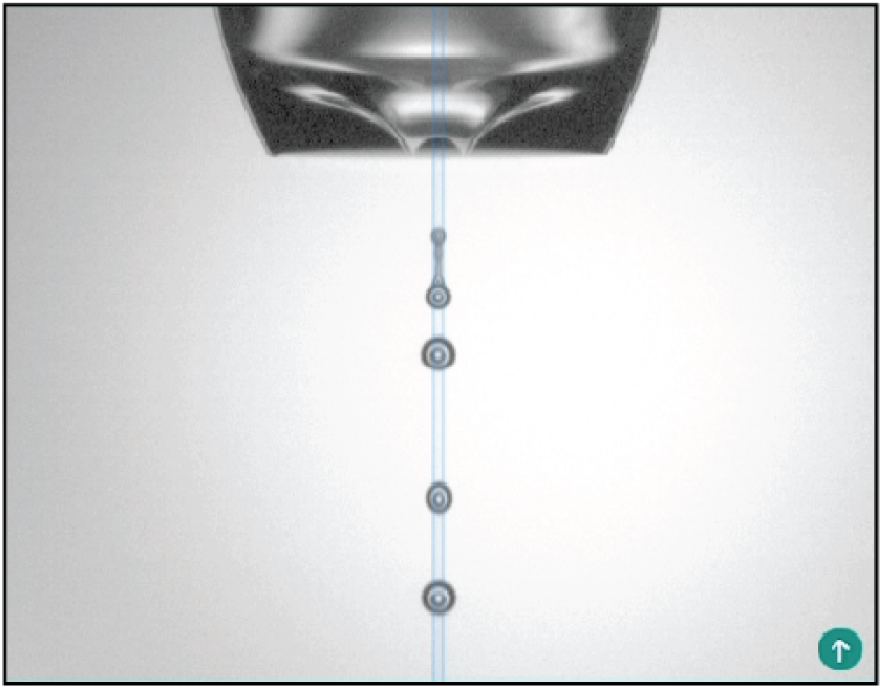
Imaged dispenser ejecting four droplets. This is the correct amplitude setting necessary for proper sample deposition. If this does not occur, refer to **Section G.2** to diagnose and fix dispenser issues
  A.17.2 **NOTE:** The troubleshooting tab provides different modes for cleaning the tip of the dispenser. This will help with proper dispensing if the automated test fails (see *Troubleshooting* **Section G.2**).
  A.17.3 **NOTE:** The advanced tab allows the user to increase or decrease the dispenser amplitude to ensure the correct placement of the four to five droplets.
A.18 Click the “Continue” button to advance to the <u>Load Cryogens</u> page.
A.19 Open the LN_2_ drawer by clicking the “Open” button, locating it to the out position.
A.20 Place the LN_2_ Styrofoam dewar in the LN_2_ drawer with the sensors facing toward the user.
  A.20.1 **NOTE:** Gently handle the Styrofoam dewar by the edges as not to damage the sensors and ensure correct seating.
A.21 Once properly seated, click the “Load Cryogen” button. The LN_2_ drawer will partially close, and the software will detect if the LN_2_ Styrofoam dewar is loaded correctly. The LN_2_ drawer will then remain in the partially closed position.
A.22 Pour LN_2_ directly into the ethane cup and surrounding dewar until the LN_2_ level in the dewar is nearly flush with the Styrofoam edges. The temperature sensor readings in the side panel will turn red and drop rapidly from room temperature as the system cools. When the temperature reaches -173 ^*°*^C, the text will turn white, and the liquid ethane can be prepared.
  A.22.1 **NOTE:** Do not rest the weight of the LN_2_ dewar on the drawer as to prevent damage to the LN_2_ drawer and/or the Styrofoam dewar.
  A.22.2 **NOTE:** The temperature sensor for the LN_2_ dewar is measured through a thermocouple attached to the ethane cup. The software maintains the ethane cup temperature at -175^*°*^C by default, although this value can be changed on the <u>Load Cryogens</u> page in the input box located below the “Close” button.
  A.22.3 **NOTE:** A warning may appear that reads *“Ethane Heater temperature out of range. Target temperature -175*.*0*^*°*^*C*.*”* This can be ignored and closed. It will not appear again once the temperature is in the range of -175^*°*^C.
  A.22.4 **CAUTION:** LN_2_ is a cryogen that can pose a serious threat to life and/or injury if not handled properly. Ensure all personal protective equipment is utilized to minimize risk of injury. The vapor of LN_2_ is an asphyxiant and should be handled in well-ventilated areas. Please consult an expert if unsure how to operate or handle cryogenic vessels and cryogens. Please refer to the institution’s Environmental Health and Safety guidelines when handling cryogens.
A.23 Open the ethane tank valve and adjust the regulator attached to the ethane tank until the outlet pressure reading is 35-40 psi when the ethane button is depressed.
  A.23.1 **NOTE:** The pressure reading when the dispenser button is depressed is critical as this establishes proper ethane gas flow.
  A.23.2 **NOTE:** If the ethane tank pressure is not set to 35-40 psi, then the time it takes to reach the proper ethane liquid level in the cup will vary greatly.
  A.23.3 **NOTE:** Carefully handle the ethane dispenser only by the Styrofoam layer to prevent damage to the dispenser.
  A.23.4 **CAUTION:** Compressed ethane gas is flammable and can pose a serious threat to life and/or result in injury if improperly handled. Please consult an expert if unsure how to operate or handle compressed gas tanks. Please refer to the institution’s Environmental Health and Safety guidelines when handling flammable compressed gases.
  A.23.5 **CAUTION:** Liquid ethane is a cryogen that can pose a serious threat to life and/or injury if not handled properly. Ensure all personal protective equipment is utilized to minimize risk of injury. Please consult an expert if unsure how to operate or handle cryogenic vessels and cryogens. Please refer to the institution’s Environmental Health and Safety guidelines when handling cryogens.
A.24 Once the LN_2_ dewar reads -175^*°*^C and the ethane dewar is devoid of LN_2_, place the chameleon ethane gas dispenser flush with the ethane cup (**Figure 3**).

**Figure 3:**
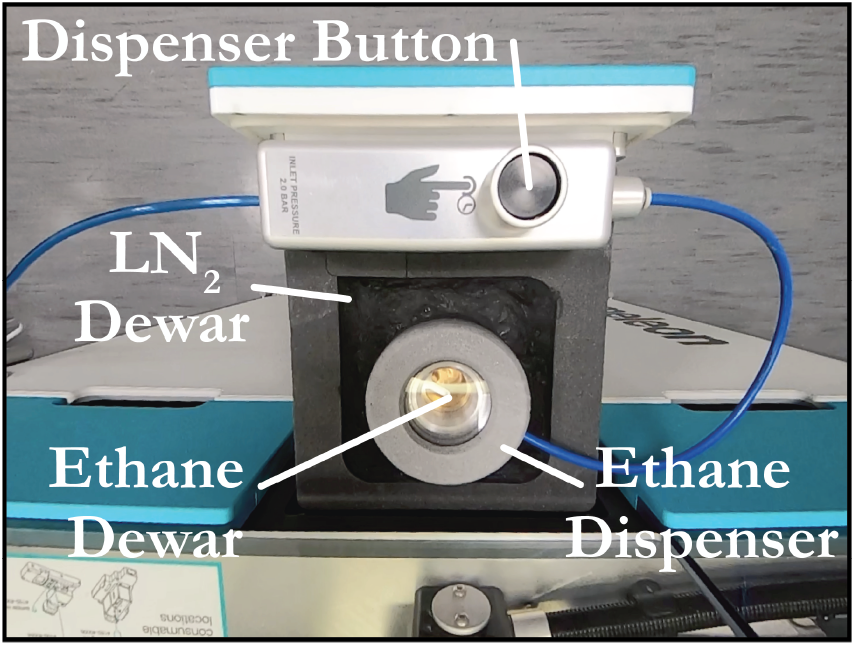
Chameleon ethane gas dispenser. The LN_2_ Styrofoam dewar and chameleon gas dispenser with LN_2_ present. Areas of interest are labeled.
  A.24.1 **NOTE:** Once the ethane gas dispenser is placed on the ethane cup, the temperature sensor will read a higher temperature and the software will turn the ethane cup heater off to allow re-cooling. Once the temperature reaches -175^*°*^C again, the software will turn the heater on and the ethane cup temperature of -175^*°*^C will be maintained.
A.25 Depress and hold the ethane dispenser button to allow ethane gas flow into the ethane dewar. Hold the button for *∼*45-48 s until the liquid ethane level reaches 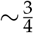 full.
  A.25.1 **NOTE:** If the ethane tank pressure is not set to 35-40 psi, then the time it takes to reach the proper ethane liquid level in the cup will vary greatly.
  A.25.2 **NOTE:** If the button is depressed too long the liquid ethane will contact the ethane gas dispenser nozzle and potentially freeze it to the ethane vessel. Great care should be taken to displace the ethane gas dispenser nozzle if this occurs.
  A.25.3 **NOTE:** If the liquid ethane level is too low, the EM grids will not be fully immersed in liquid ethane, preventing proper vitrification of the sample and potentially loss of sample/grid.
A.26 Once the optimal liquid ethane level has been reached, carefully remove the ethane gas dispenser from the ethane cup and close.
  A.26.1 **NOTE:** If the ethane dispenser remains on the cup too long or ethane splashes onto the dispenser after ethane dispensing (i.e., 5-10 s), it can freeze to the ethane cup. See *Troubleshooting* **Section G.7** for instructions to remove ethane dispenser and recover LN_2_ box.
  A.26.2 **NOTE:** Carefully handle the ethane dispenser only by the Styrofoam layer to prevent damage to the dispenser.
  A.26.3 **NOTE:** If the LN_2_ level has dropped below the Styrofoam edges then add more LN_2_ before continuing.
A.27 Click the “Close” button to close the drawer then click the “Continue” button to advance to the <u>Replace Grid Button</u> page.
A.28 Enter the ID of the white grid box.
  A.28.1 **NOTE:** White grid boxes are called “buttons” in the chameleon® software.
  A.28.2 **NOTE:** If the user forgets to add a white grid box while the LN_2_ drawer is open in **Section A.19-A.26**, the “Open” button on the <u>Replace Grid Button</u> page will re-open the LN_2_ drawer and allow the user to add a white grid box.
A.29 Click the “Continue” button to initiate tweezer cleaning. This will dip the tweezers in ethanol and allow them to air dry.
A.30 The software will automatically advance to the Load Grids page.

### B Initialize the chameleon and Prepare Grids for Sample Vitrification

This section outlines the procedure for loading grids, optimizing dispenser parameters, and preparing samples with an estimated operational time of 5-10 mins. Adjustments to the dispenser’s settings are necessary, particularly for viscous samples, to achieve the desired dispensing properties before proceeding to plunge control. Recommendations for glow discharge duration and current are provided to optimize sample wicking and deposition properties. This process is critical to ensure that the grids are properly prepared, and the sample is accurately deposited, which is vital for successful sample vitrification.

B.1 After the software has advanced to the <u>Load Grids</u> page, the lights will turn green and the doors will unlock. Using clean tweezers, carefully place the desired number of grids in the grid holder, starting from position I located in the bottom left, moving away from the door (**Figure 4**). Close the doors when finished.

**Figure 4:**
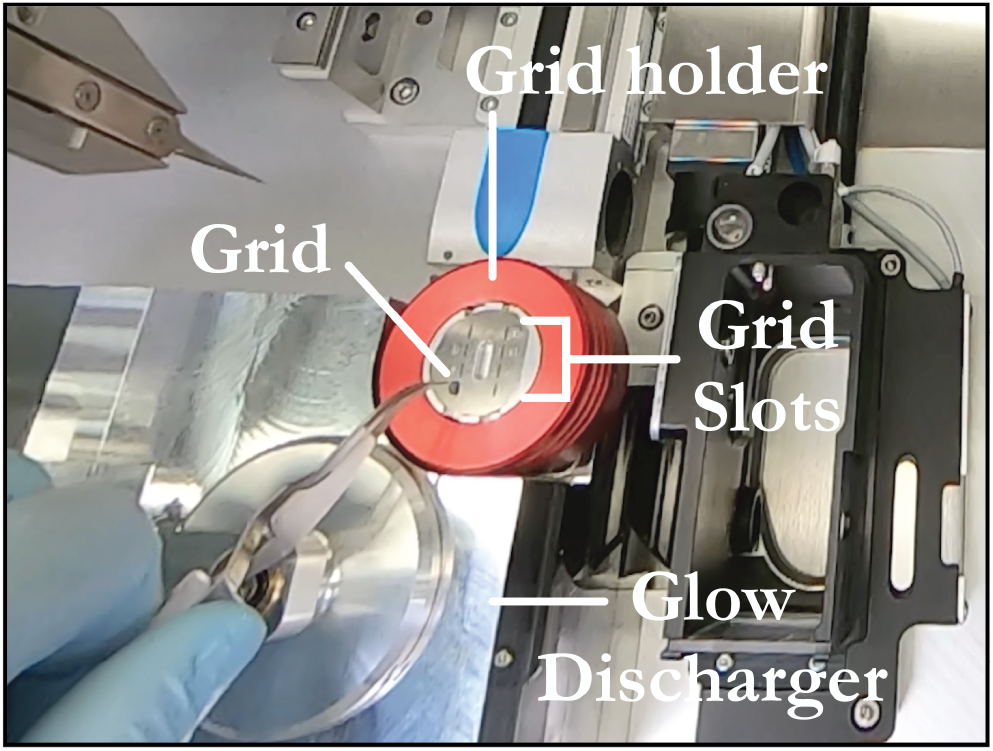
Grid placement within the chameleon. The grids should be loaded with the iridescent side of the grids facing towards the left of the instrument. The chameleon can be loaded with 8 grids at a time. Areas of interest are labeled.
  B.1.1 **NOTE:** Make certain that the iridescent (e.g., “rainbow”) side of the grids are located towards the left of the instrument, away from the dispensing tip, to ensure that the nanowires will be correctly located towards the dispenser for “self-wicking” post sample deposition.
  B.1.2 **NOTE:** Grids should only be handled using tweezers by the outer ring to prevent damage of the grid foil that could result in complete loss of grid squares and/or damage to the nanowires that could yield inconsistent results. Additionally, bending the grid during placement can lead to inconsistent behavior during glow discharge, sample deposition, and/or sample wicking.
  B.1.3 **NOTE:** The chameleon grid holder can support up to 8 grids. Grids should always be loaded starting at position I since that is the default loading position for the software.
B.2 Enter the grid batch number and grid box position information (e.g., grid box column and position number) in the *Load Grids* panel
B.3 Set the glow discharge time in the “Duration” input box located in the *Grid Activation* panel.
  B.3.1 **NOTE:** It is possible to change the current applied to the glow discharge unit, but it is recommended to leave the current input at the default value (e.g. 12 mA).
B.4 Set the maximum number of grids to load in the glow discharge unit in the “What’s the maximum number of grids to load into the glow discharge unit?” input box located in the *Grid Activation* panel.
  B.4.1 **NOTE:** A maximum of 4 grids can be glow discharged concurrently.
  B.4.2 **NOTE:** The minimum time allowed is 1 s and the maximum is 1800 s. This parameter should be tuned based on the desired wicking properties and sample viscosity. 20 - 40 s is recommended.
  B.4.3 **NOTE:** When glow discharging four grids at a time, we have noticed inconsistent wicking behavior between the first and fourth grids when using shorter glow discharge times (e.g., 20-40 s), presumably, due to the time delay between freezing each grid resulting in changes in grid hydrophilicity. If the glow-discharge time is longer (e.g., 80-120 s), this phenomenon is not as prevalent but still observable. For consistent wicking results, the user may want to glow discharge one grid at a time.
B.5 Click the “Continue” button. The robotic tweezers will grab the first grid (starting from position I) and place it in the glow discharge unit. If more than one grid is requested, then each grid will also be sequentially placed in the glow discharge unit (4 grids maximum). The glow discharge unit lid will close, and vacuum will be pulled. The glow discharger runs for the requested duration.
  B.5.1 **NOTE:** Once this step is initiated it cannot be interrupted.
  B.5.2 **NOTE:** The user can verify that the glow discharge unit is functioning by observation of a purple plasma within the glow discharge unit. The software will automatically advance to the <u>Prepare Sample</u> page.
B.6 Once glow discharging is complete, the doors will unlock, and the monitoring lights will turn green. Enter the sample block temperature in the input box located in the *Load Sample* panel, “Sample block temperature set point”.
  B.6.1 **NOTE:** The sample block temperature is set to 4.0^*°*^C by default.
B.7 Enter the desired sample volume that will be aspirated into the dispenser using the “Volume to aspirate” input box located in the *Load Sample* panel. The minimum volume permitted to load into the dispenser tip is 3 *µ*L which requires a minimum of 5 *µ*L to be loaded into the sample vial to ensure no air enters the dispenser tip during aspiration.
B.8 Enter the sample details in the *Sample Details* panel. The information for this sample will be saved and the user will not need to re-enter it in the future.
B.9 Carefully pipette the sample whilst ensuring no air bubbles are introduced into the sample.
  B.9.1 **OTE:** It is recommended that an additional 2-4 *µ*L of sample above what will be aspirated is loaded into the sample vial to ensure proper aspiration and prevent introduction of air bubbles into the dispenser tip.
  B.9.2 **NOTE:** We recommend aspirating 8-10 *µ*L of sample for grid preparation with 12-15 *µ*L of sample loaded into the vial. This will allow enough sample volume for dispenser troubleshooting procedures that may be required.
  B.9.3 **NOTE:** For highly viscous samples, ensure the meniscus is as level as possible to prevent uneven sample draw and the introduction of air bubbles.
  B.9.4 **NOTE:** : Since the dispenser tip loads sample from the bottom of the sample vial, no debris or air bubbles should be present otherwise loss of sample or damage to the dispenser tip may occur.
B.10 Click the “Continue” button to initiate sample aspiration. The software will automatically advance to the <u>Retest Dispenser</u> page when finished.
B.11 Click the “Dispense” button to test the dispenser. An image of the dispenser tip and the sample droplets exiting the dispenser tip will appear.
  B.11.1 **NOTE:** : In **NOTE A.17.1** of **Section A.17**, we recommended four to five equally spaced droplets for ideal sample deposition. The dispenser test at that step was for pure water. To obtain the ideal four to five droplets with a protein sample the amplitude of the dispenser will need to be adjusted and/or the tip will need to be cleaned since the sample will most likely be more viscous than pure water. See **Section G.1** for troubleshooting dispenser amplitude and issues.
  B.11.2 **NOTE:** The user should not proceed until the desired dispensing properties are attained otherwise suboptimal grids will result.
B.12 Click the “Continue” button to advance to the <u>Plunge Control</u> page.

### C Evaluate Sample Deposition and Tune Plunge Times for Optimal Sample Deposition and Wicking

This section details the process for evaluating sample deposition and optimizing plunge times to achieve optimal sample deposition and wicking, with an estimated operational time of 30-40 mins. Users can manually alter both plunge and wicking times to optimize sample deposition for their specific sample. There are two modes for sample deposition. The two-stripe mode is primarily used for initial analysis and is recommended for the first few grids of a new batch or sample. If results are satisfactory, one-stripe mode is used for subsequent grids. The software allows the user to review the deposition results via video. The user can either accept or reject the grid based on comparison with standard wicking results. Accepted grids are stored in the supplied grid box, and rejected ones are discarded.

C.1 By default, the two-stripe dispensing mode will be automatically selected. In the *Test Wicking* panel, click the “Characterize” button. One stripe is deposited offset from the grid center and a video of the sample wicking is presented to the user for evaluation. This allows the user to evaluate sample application and the wicking time. The software will recommend a plunge time for the second dispensed stripe. Click the “Plunge” button to initiate the second dispense and plunge freeze (**Figure 5**).

**Figure 5:**
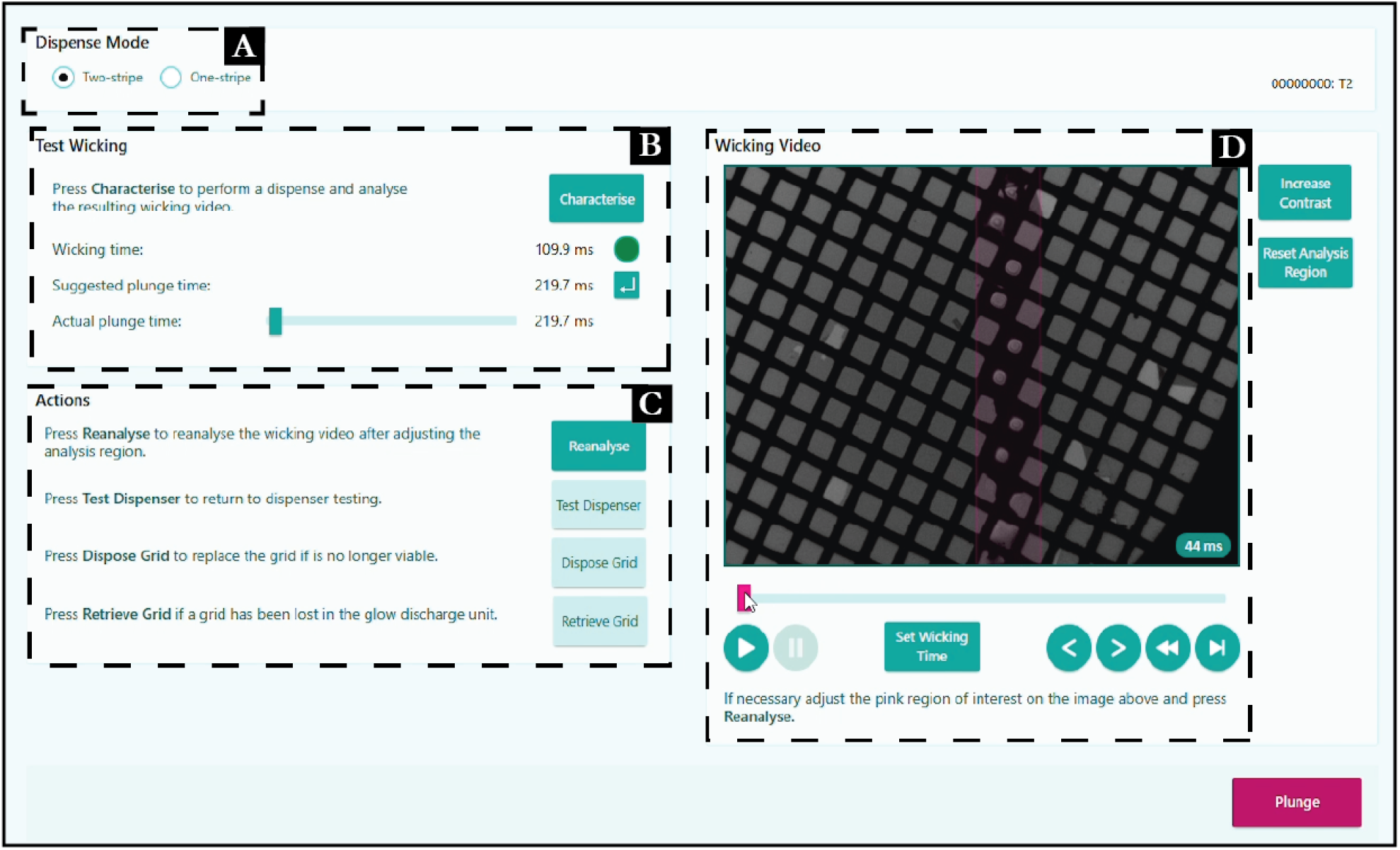
Two Stripe mode page with highlighted areas. A) Dispense mode selection panel. B) Test Wicking panel. Recommended wicking times can be found within this area. C) Actions Panel. D) Wicking Video which displays the results of the two stripe mode
  C.1.1 **NOTE:** In this mode, the grid is not vitrified if the “Characterize” button is clicked but retained for the second stripe. At the end of deposition of the second stripe, the sample will be vitrified.
  C.1.2 **NOTE:** If the characterization grid in two-stripe mode is used for plunge freezing, the <u>Plunge Review</u> page will appear after clicking the “Plunge” button on the <u>Plunge Control</u> page in two-stripe mode. See **Section C.3**.
  C.1.3 **NOTE:** If the user is not satisfied with the software recommended plunge time or wicking time, the user can manually adjust either the plunge time using the “Actual plunge time” slider bar in the *Test Wicking* panel or the wicking time using the slider under the video in the *Wicking Video* panel and clicking the “Set Wicking Time” button under the slider.
  C.1.4 **NOTE:** Keep in mind that two-stripe mode is for initial analysis of grid and sample properties. Users should use the wicking time from two-stripe as a starting point for dialing in the plunge time in one-stripe mode.
  C.1.5 **NOTE:** We recommend using two-stripe mode for the first 1-2 grids from a new batch of grids and/or the first time a sample is being prepared. Therefore, we do not recommend this for every freezing session unless so desired by the user.
  C.1.6 **NOTE:** The characterization grid can be used for data collection, although the quality of the grid and data collection results will vary from single stripe mode grids (see **NOTE C.1.7**). If the user wants to keep this grid for collection, follow instructions in **Section C.3-C.9**.
  C.1.7 **NOTE:** It has been observed that solution from the first stripe can fenestrate across the nanowires to squares 2-4 rows away from the column. This can alter the wicking properties of the second stripe leading to inconsistent behavior (i.e., thicker ice) and therefore should be primarily only used for diagnostic purposes.
C.2 If the user is satisfied with the two-stripe mode results, one-stripe mode should be used for the rest of the grids. In this setting, the user can manually define the plunge time and click the “Plunge” button to initiate sample freezing.
  C.2.1 **NOTE:** It is recommended to click the “Test Dispenser” button and confirm the four droplets before each dispense and plunge freeze. Air bubbles may be introduced into the dispenser when inspecting and testing the dispenser. See **Section G.3** to troubleshoot air bubbles.
C.3 The software will automatically advance to the <u>Plunge Review</u> page. In the *Review Dispense Video* panel, a video will show the wicking results. The grid can either be accepted or rejected. If the grid is accepted, the grid will be placed into the supplied grid box, else the grid is discarded (**Figure 6**).

**Figure 6:**
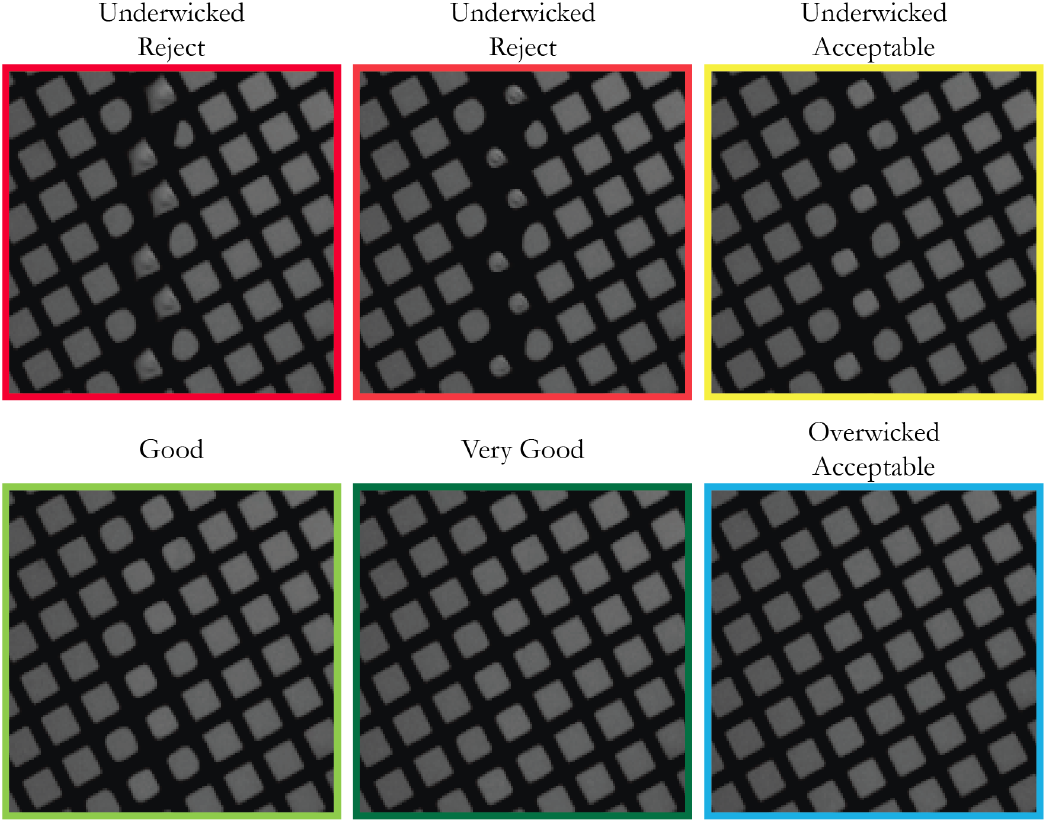
Reference images for different wicking times. These serve as a guide for determining proper wicking times for the creation of suitable grids for data collection. Adapted from SPT labtech.
  C.3.1 **NOTE:** Six images are shown below the dispense video. They represent examples of different wicking results for comparison to the wicking result in the review video. It is recommended that the user’s result match closely to the Very Good or Overwicked Acceptable comparison images.
  C.3.2 **NOTE:** Ensure the tweezers within the LN_2_ have released the grid into the grid box. If the grid remains stuck to the tweezers, click the “Retry” button.
C.4 Click the “Lift Tweezers” button when the grid has been properly stored.
C.5 Click the “Continue” button to initiate the automatic tweezer cleaning process. During this process, the <u>Accept Grid</u> page will appear and allow the user to fill in comments and select the appropriate option to describe the grid.
C.6 Click the “Continue” button to advance to the <u>Another Grid?</u> Page.
C.7 If more grids are to be frozen, click the “Prepare Grid” button to advance to the <u>Reload Glow Discharge</u> page. This will allow additional grids to be glow discharged and frozen. If the user is finished, proceed to **Section D.1**.
C.8 If the user is glow discharging one grid at a time, then confirm the “Perform Additional Glow Discharge” box is checked and enter the glow discharge duration. Enter the maximum number of grids to place in the glow discharge unit. Click the “Continue” button to initiate the glow discharge process.
  C.8.1 **OPTIONAL:** It is recommended to run an additional air dry on the tweezers between plunge freezing and glow discharging more grids. This can help reduce grids sticking to the tweezers. Click the “Dry Tweezers” button on the <u>Reload Glow Discharge</u> page.
  C.8.2 **NOTE:** If the user initially and simultaneously glow discharged more than one grid (e.g. up to 4 grids), then the “Perform Additional Glow Discharge” box is automatically unchecked and cannot be selected.
C.9 Click the “Continue” button to advance to the <u>Plunge Control</u> page where the user will repeat **Sections C.1-C.8**.

### D Finish the Session and Clean the chameleon

This section outlines the procedures for concluding a session and cleaning the chameleon system, with an estimated operational time of 10-15 mins. The dispenser cleaning process includes a thorough automatic test to ensure the dispenser’s functionality. If the session involves continuing with the same or a new sample, appropriate steps are followed to replace or retain the white grid box using the software prompts, ensuring proper identification and placement of grid boxes. If no new samples are needed, the process moves to preparing for system shutdown, including the removal of unused grids. The user is given the option to create detailed reports and save sample deposition and wicking videos, which document the inputs and results for future reference. Proper dispenser cleaning and system shutdown is important to maintain consistent sample deposition and wicking results.

D.1 If the user is finished freezing grids or wants to load a new sample, then click the “Finish Session” button on the <u>Another Grid?</u> page to initiate the dispenser cleaning process. This will take 7-10 mins.
  D.1.1 **NOTE:** Once this process is initiated, the sample in the dispenser will be lost. This process cannot be interrupted.
D.2 The software automatically advances to the <u>Test Dispenser</u> page. Click the “Automatic Test” button to initiate the automatic dispenser test. If the dispenser test passes, the user should see a “Dispenser tested ok” message.
D.3 If the user needs to load a new sample, click the “Load Sample” button. This will take the user to the <u>Replace Grid Button</u> page. Otherwise go to **Section D.10**.
D.4 At this stage, the user may continue using the same white grid box if there are slots available. Click the “Keep Grid Button” button to keep the same white grid box.
D.5 If the user does not want to mix samples in the same white grid box, a new one can be added. Click the “Open Drawer” button to open the LN_2_ drawer. The user can move the current white grid box to one of the three Styrofoam slots on the left side of the Styrofoam dewar and place the new white grid box in the white grid box collection slot.
D.6 Enter the white grid box ID in the “Enter the ID of the new button” input box.
D.7 Click the “Close Door” button.
D.8 Click the “Continue” button to advance to the <u>Load Grids</u> page.
D.9 Follow the instructions in **Section B.1-B.12**.
D.10 However, if the user does not need to load a new sample, then click the “Continue” button to advance to the <u>Dry Dispenser</u> page.
  D.10.1 **NOTE:** If there are unused grids in the glow discharge unit, the robotic tweezers will remove them from the glow discharge unit and place them back in their grid holder positions.
D.11 The lights will turn green, and the doors will unlock. Refill the methanol vial. Close the doors.
D.12 Click the “Dry” button to initiate dispenser cleaning and drying with methanol. The software automatically advances to the <u>Remove Cryogens</u> page.
  D.12.1 **NOTE:** If there are unused grids in the grid holder, the software will advance to the <u>Remove Grids</u> page first. The grids can be recovered and placed in the user’s grid box. When done, click the “Continue” button to advance to the <u>Remove Cryogens</u> page.
D.13 Click the “Open” button to open the LN_2_ drawer. The user should move the white grid box(s) to a separate dewar filled with LN_2_.
D.14 The Styrofoam LN_2_ dewar can be removed from the drawer slot and the LN_2_/ethane safely disposed of. Do not place the Styrofoam dewar back in the chameleon. Click the “Close” button to close the LN_2_ drawer. Click the “Continue” button to advance to the <u>Finished</u> page.
D.15 Click the “Create Reports” button to advance to the <u>Report</u> page. Ensure “All grids, by session” is highlighted in the list of options on the left. Select the most recent session in the “Choose session” list. Click the “Create Report” button at the lower right of the page. The user can save the report to a designated folder.
  D.15.1 **NOTE:** This report includes information about the inputs and results from the session. This should be used for future grid freezing parameter decisions.
D.16 On the Report page again, highlight the “Videos Only” option on the left of the page. Choose the most recent session from the “Choose session” list. Click the “Save All” button. Save the session videos to a designated folder.
  D.16.1 **NOTE:** The videos show the dispenser and wicking results. This should be used for future grid freezing parameter decisions.
D.17 Click the grid symbol at the top of the page on the right. This will return to the <u>Finished</u> page. Click the “Close” button. The chameleon light will turn blue and the chameleon® software will close. The doors are unlocked when the light is blue.
D.18 Remove the sample vial and dispose of it.
D.19 The humidifier bottle should be removed from the humidifier, refilled (See **Section A.5**), and stored in a space next to the humidifier.
D.20 Empty the used wash fluid bottle (see **Section A.3**).
  D.20.1 **NOTE:** Methanol is collected in this bottle. Consult the institution’s Environmental Health and Safety guidelines when handling and disposing of methanol.
D.21 The wash pump supply bottle should be refilled with ultrapure water (see **Section 1.2**).
D.22 The chameleon session is now complete.

### E Grid Evaluation in the Transmission Electron Microscope

This is a general guide for high-throughput screening and data collection from grids prepared using the chameleon. Estimated operating time: 10 mins - 2 hrs. This section assumes the user is familiar with the operation of a latest generation transmission electron microscope (TEM) and has been trained to load samples and operate a TEM independently. The ensuing sections will provide a general workflow using a ThermoFisher Titan Krios G4 equipped with a Falcon 4 direct electron detector with Selectris-X energy filter operated using ThermoFisher Tecnai User Interface (TUI) and EPU 2.1+ software packages. The general principles provided below can be adapted to other TEMs but will have to be modified depending on software available, desired magnification and exposure rate, and user preferences.

E.1 Follow the recommended guidelines for clipping grids into thermofisher scientific autoGrids for loading into the TEM. Please refer to **[cryoEM merit badge: Grid Clipping](https://www.cryoemcenters.org/merit-badge/tfs-autogrid-clipping/)** for additional details.
E.2 Load the grid(s) into the microscope following recommended procedures. Inventory the grids by clicking on the “Inventory” button in the Autoloader tab in TUI. Please refer to **[cryoEM merit badge:Autoloader](https://www.cryoemcenters.org/merit-badge/tfs-autoloaders/)** for additional details.
E.3 Navigate to the “Atlas” tab in EPU and select the “Session Setup” option in the *Task* panel. Click the “New Session” button in the top left and enter the appropriate information in the *Edit Session* panel. Please refer to **[cryoEM merit badge: Screening with EPU](https://www.cryoemcenters.org/merit-badge/screening-with-epu/)** for additional details.
E.4 Select the grid position(s) to be atlas’ed by clicking the empty box next to the slot number. Once all grids have been selected hit the “Start” button in the top left.
E.5 The software will then proceed to acquire low magnification images of each EM grid that will be stitched together in a composite image termed an Atlas. This is necessary to select grid squares for downstream screening and data collection.
  E.5.1 **NOTE:** This process will generally take 10 mins per grid.
E.6 After all the atlases have been acquired, examine the Atlas image of each grid for the presence of the ice stripe. Most chameleon-prepared grids will have empty squares so the sample deposition stripe can usually be identified by finding contiguous darker grid squares. However, areas of thin ice can potentially be harder to identify due to the low contrast set by default in EPU.
  E.6.1 **NOTE:** To help determine the sample stripe, one can adjust the histogram slider in the lower right panel to increase contrast.
E.7 The ice stripe can be identified by the difference in contrast from surrounding empty grid squares. For areas with the thinnest ice possible, the contrast between empty grid squares and grid squares with ice is minimal (**Figure 7**). To confirm the presence of ice, it may be necessary to load the grid, if currently not on the stage, and take diagnostic images of the grid squares of interest. To load the grid, click the “Load Sample” button found in the top panel. Once the grid is loaded, identify grid squares of interest, and take a “Hole/Eucentric Height” image. This will allow the user to accurately identify empty and filled holes based on the intensity of the holes, with brighter being empty.

**Figure 7:**
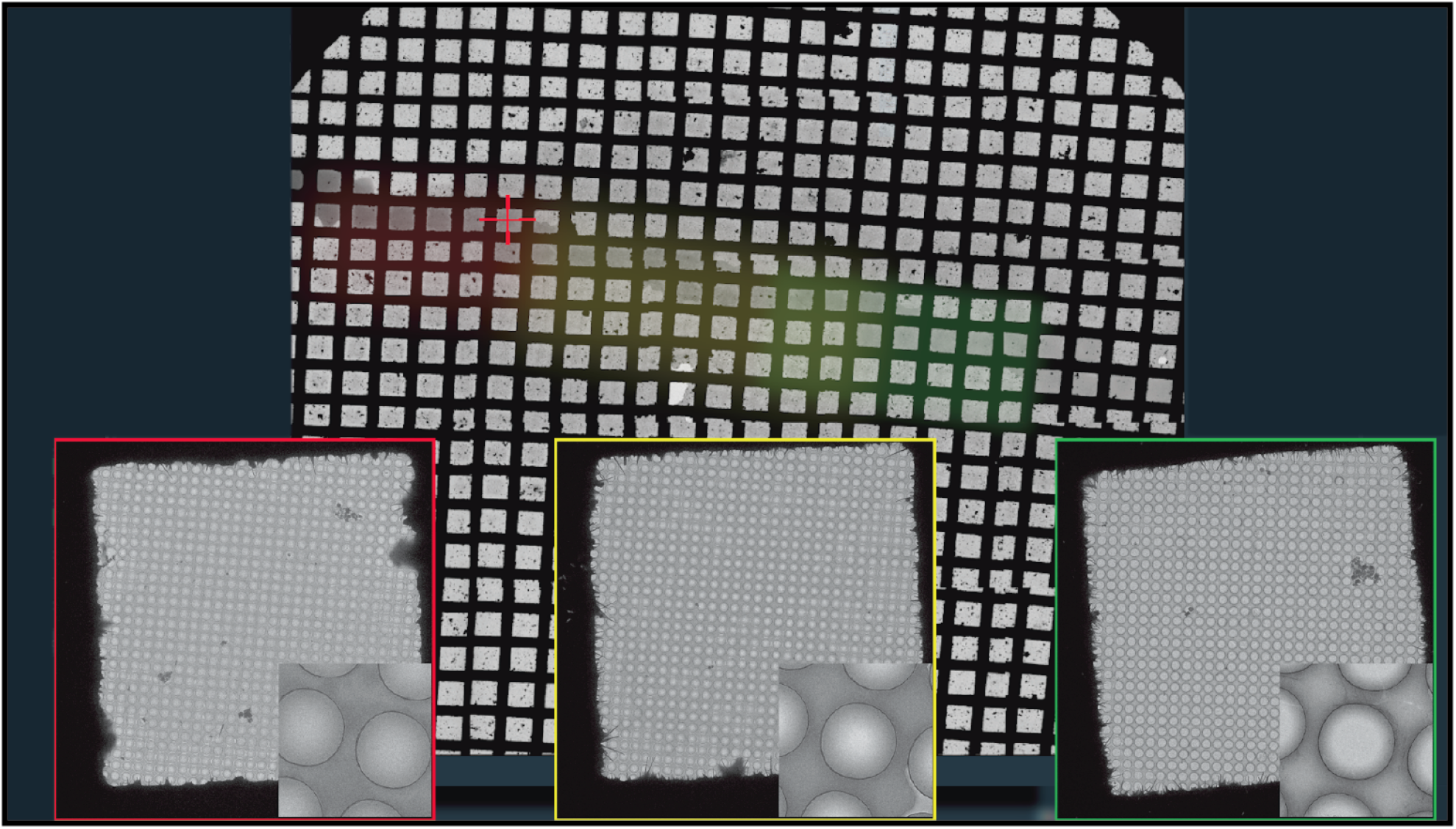
A representative atlas image displaying the ice thickness gradient along the ice stripe. Squares from the thick region (red), the moderately thin area (yellow) and desirable thin area(green) are displayed bellow.

**Figure 8:**
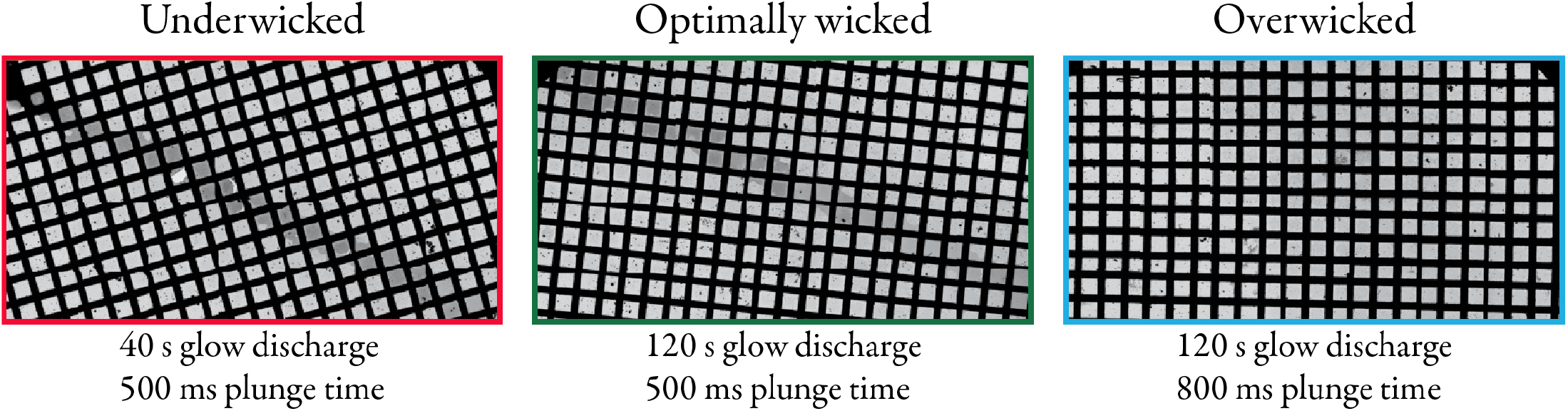
Demonstration of the influence of glow discharge and plunge times on ice stripe quality. Increasing the glow discharge time while keeping the plunge time the same yielded desired ice thickness (left to middle). Further increasing the plunge time yielded a grid that was overwicked with limited real estate for collection (middle to right).
  E.7.1 **NOTE:** This will load the selected grid and correctly reposition the Atlas image of that square for any rotation/translation errors that resulted from grid unloading and reloading. The process of switching grids will take approximately 4 mins.
  E.7.2 **NOTE:** If the Atlas of the last grid that was acquired is deemed the “best”, that grid will still be loaded in the microscope’s CompuStage and loading will not be required.
  E.7.3 **NOTE:** Depending on the nature of the sample, screening multiple ice thicknesses may be necessary to find particles that lead to a useful reconstruction.
E.8 Once grid squares with ice thickness appropriate for the sample have been identified, it is recommended to take diagnostic images of a few holes from each grid square to ensure the presence of particles and distribution of the sample within the ice. If particle density and distribution is uniform across the micrograph, it is recommended to continue on to high-throughput data collection.

### F Ice Thickness Tuning by Glow Discharge and Plunge Time Optimization

Following grid evaluation, if the ice thickness is determined to be too thick and incompatible with high-quality image recording, we recommend that the user optimize the plunge time and/or glow discharge time. By fine-tuning the plunge time and glow discharge time, users can effectively tune the ice thickness and particle density. Below, we outline recommendations for adjusting these parameters that can help to achieve the desired ice thickness for optimal results. Of note, this assumes that particle density and particle distribution are adequate, but thinner ice is required for high-quality imaging.

#### F.1 Tuning plunge and glow discharge times

F.1.1 The glow discharge duration is set to 20 s by default, but users have the flexibility to adjust this value within the range of 1-1800 s. If the user desires a shorter plunge time than the recommended range of 400-600 ms but still aims for very good to overwicked grids, compensating by increasing the glow discharge duration is necessary.
F.1.2 If a wider plunge time range is desired while maintaining very good to overwicked grids, the glow discharge duration should be adjusted accordingly.
F.1.3 There is an available plunge time mode of 54 ms; however, this mode does not generate a review video for wicking analysis and can yield inconsistent results in sample deposition.
F.1.4 If users require a faster plunge time but seek more consistent outcomes than the 54 ms mode, it is recommended to increase the glow discharge duration to 120-150 s. Beyond this duration, the wicking process becomes very rapid (almost instantaneous).
  F.1.4.1 Setting the plunge time to 100-150 ms will provide a review for wicking analysis and can result in grids ranging from very good to overwicked while maintaining a brief plunge time.
F.1.5 We recommend adjusting glow discharge time in 5-10 s intervals while keeping the plunge time the same. Conversely, we recommend adjusting the plunge time in 50 ms intervals while keeping the glow discharge the same.

#### F.2 Tuning plunge and glow discharge times for low-viscosity samples

F.2.1 For low-viscosity samples (e.g., samples lacking glycerol or detergents), we have determined that a glow discharge duration between 40-60 s will ensure adequate wicking and sample thinning when combined with a plunge time to be set between 400-600 ms. These ranges of values typically result in very good or overwicked grids.

#### F.3 Tuning plunge and glow discharge times for high-viscosity samples

F.3.1 While the default setting works well for most samples, protein samples at high concentrations, glycerol or other polyols, or detergents/surfactants may require a longer glow discharge duration. In the case of high-viscosity samples with 5-10% glycerol or detergents, extending the glow discharge duration to 80-120 s will maintain the plunge time within the 400-600 ms range while producing very good or overwicked grids.

### G Troubleshooting

#### G.1 Sample is not dispensing properly

The default amplitude for the dispenser was obtained against pure water and therefore will most likely not match the viscosity of your sample.

G.1.1 If less than three drops are observed, then navigate to the <u>Troubleshooting Tab</u> and increase the amplitude. We recommend working in increments of 25. Conversely, if more than four droplets are observed, then decrease the amplitude.
  G.1.1.1 **NOTE:** The <u>Troubleshooting Tab</u> is found on the <u>Retest Dispenser</u> page following sample aspiration on the <u>Prepare Sample</u> page (**Section B.10**) or on the <u>Plunge Control</u> page by clicking the “Test Dispenser” button.
G1.2 If no drops are observed then there could be debris, an air bubble, or insufficient back pressure in the dispenser (see **Section G.2 or G.3**).

#### G.2 Debris in or on dispenser

The tip of the dispenser can attract debris and must be cleared for proper sample dispensing. The debris often comes from the wiping pad but can be cleared using four options in the <u>Troubleshooting Tab</u>. Starting with **Section G.2.1.1** and proceeding to **Section G.2.1.2-4** as needed will clear most debris blocking the dispenser.

G.2.1 The “Wipe Dispenser” button will perform one dispenser wipe on the pad.
G.2.2 The “Multi-Wipe Dispenser” button will perform multiple dispenser wipes on the pad.
  G.2.2.1 **NOTE:** Wiping the dispenser usually clears the debris, but it can increase the debris on the dispenser tip.
G.2.3 The “Dip Dispenser” button dips the dispenser in the washing station. This works if the debris is soluble or small.
G.2.4 The “Mini Prime” button dispenses 0.5 *µ*L of solution. If the other three methods don’t work, this can be used to clear larger debris.

#### G.3 Air bubble in the dispenser tip or insufficient back pressure

Air bubbles can develop in the dispenser tip from aspiration of sample or other causes. This can impede the dispenser flow causing inconsistent sample deposition.

G.3.1 The dispensing action may dislodge the air bubble(s). This can be accomplished by repeatedly clicking the “Dispense” button.
G.3.2 If **Section G.3.1** is unsuccessful, there are three additional options in the <u>Troubleshooting Tab</u>:
  G.3.2.1 The “Mini Prime” button dispenses 0.5 *µ*L of sample through the dispenser. The larger volume of sample pushed through the dispenser tip can dislodge the air bubble.
  G.3.2.2 In the *Advanced* panel, the user can change the “Amplitude”. Increasing the amplitude from the default value increases the strength of dispensing. Increasing this value may dislodge the air bubble.
  G.3.2.3 The “Jog Bottle Up” and “Jog Bottle Down” buttons change the height of the dispenser bottle in increments of 0.5 mm. This changes the back pressure in the dispenser which may dislodge the air bubble. This can have greater success when used in conjunction with changing the amplitude in **Section G.3.2.2**.

#### G.4 No stripe detected

After sample deposition (see **Section C.3**), the review video will show sample wicking. Usually, the stripe is observable in the first frames of the video. If no stripe is observed, this could mean the dispenser did not deposit sample onto the grid. It is recommended to test the dispenser before each sample deposition. If the dispenser is working as expected, then the wicking may have occurred quickly, and no stripe is observed.

G.4.1 **NOTE:** If the wicking does occur fast, then the sample is there and won’t be observable until screening the grid in the TEM.
G.4.2 **NOTE:** If a longer glow discharge and long plunge time is used (i.e. 60 s+ glow discharge and 500 ms+ plunge time), then wicking may occur very quickly and is to be expected.
G.4.3 **NOTE:** If the user selects the 54 ms plunge time option, then a review video is not provided. The stripe will not be observed until grid screening on the microscope.

#### G.5 Ethane dispenser freezing to LN_2_ box

G.5.1 Remove the LN_2_ box from the drawer and warm the dispenser using the ambient temperature of the room. Depending on the time required, the LN_2_ and ethane may still be usable if and can be placed again in the chameleon drawer slot.

## III Representative Results

Lyophilized human hemoglobin was resuspended in phosphate buffered saline (PBS) at a concentration of 5 mg/mL. Grids were prepared using the protocol described above and imaged using a ThermoFisher Scientific Titan Krios (G4) with a Falcon 4 direct electron detector and a Selectris X energy filter (165 kx magnification, 0.735 Å/pixel, 60 e-/Å^2^ cumulative exposure). All images were processed using cryoSPARC live version 4.2.1. Across a single high-quality grid, representative images were collected, and motion corrected to indicate the contribution of ice thickness to the resulting particle density, and quality, as well as the observed particle contrast. For the micrograph obtained from the thick area of the sample stripe, the particles are observed to be clumping and low contrast. The contribution of the thicker ice can be observed in the power spectrum of the thick ice micrograph as a diffuse ice ring and lower CTF estimation resolution fits (5.8 Å resolution). Images collected in thinner areas display better particle distribution, higher contrast, and better resolution in the CTF fits (∼3.1 Å for medium thick ice and ∼2.9 Å for thin ice). Notably, images acquired from the thinnest ice area are most desirable and can be obtained by imaging at the thinnest section of the sample ice stripe.

## IV Discussion

The preparation of high-quality grids remains a critically important step for successful structure determination by cryoEM, yet issues with the air-water interface and irreproducibility between grid preparations - primarily arising from various manual aspects of the vitrification process - introduce random errors and stochasticity that decrease the success rate. Although the introduction of blot-free vitrification and the automation of critical steps of the grid preparation process – from grid glow discharging to sample application and vitrification – greatly affords an opportunity to produce EM grids with more consistent and predictable ice thickness, the process is not foolproof, and care has to be taken to produce optimal results for each unique specimen. Using the steps we have outlined herein, researchers should be able to operate the chameleon to prepare high-quality EM grids for high-throughput screening and data collection. While this method can be quite robust, several aspects of the protocol must be carefully tailored to the specimen of interest based on feedback from the chameleon software and EM grid screening. We have outlined several of these critical steps below, their importance in the grid preparation workflow, and provided recommendations on how to troubleshoot these steps.

The hydrophilicity and wicking capability of the nanowire grids, dictated by the glow discharge time as well as the grid plunge time, respectively, are critically important for tuning both the overall ice thickness as well as the shape of the ice thickness gradient across the sample stripe. It is important to optimize these parameters concurrently for each sample using the real-time feedback of the camera monitoring sample wicking to obtain high-quality grids with thin ice. However, prior to tuning these parameters we recommend first optimizing the size and shape of the sample droplets obtained from the dispenser. The calibrated amplitude should produce 4 to 5 evenly spaced droplets with a water sample, so this value is typically increased to account for the viscosity of the sample, which will vary depending on sample concentration and buffer composition. Following cleaning of the dispenser tip post sample aspiration, if fewer than 4 to 5 droplets are obtained for the sample, it is recommended to increase the amplitude by intervals of 100 (not to exceed the maximum 2047) until the desired 4 to 5 droplets are obtained. At this point the amplitude can be fine-tuned in smaller increments until 4 to 5 droplets are reproducibly observed. If, however, changing the amplitude does not work, then using the default amplitude but lifting the dispenser backflow bottle in intervals of 1-2 mm using the “Jog Bottle Up” button can increase the pressure at the dispenser, resulting in sample dispensing. Once droplets are obtained, the amplitude value can then be increased as described above.Together, bottle adjustments in conjunction with amplitude adjustments should produce 4 to 5 droplets consistently, even for viscous samples. Of note, the consistent deposition of the same sample volume, dictated by the size and shape of the droplets, is necessary for tuning grid behavior. Indeed, larger than desired sample deposition volumes can require significantly longer plunge times with more variability across grids.

Following optimization of sample dispensing, we recommend focusing next efforts on concurrently optimizing the glow discharge time and plunge time to produce the ideal ice thickness stripe. The default current for the glow discharge is 12 mA, although this can be increased (i.e., 20 mA). We prefer to keep the default current value and extend the glow discharge from the default of 20 s to longer times, usually in the range of 40-90 s, but sometimes as high as 120 s or more for very viscous and/or small specimens (e.g., <80 kDa). If the sample has lower viscosity (e.g., no glycerol and/or detergents), we recommend a glow discharge duration of 40-60 s with a plunge time of 400-600 ms. This should result in a “very good” or “overwicked” grid according to the chameleon software evaluation. If the sample is more viscous, (e.g., 5-10% glycerol and/or detergents), we have found success increasing the glow discharge time to 80-120 s to allow for the plunge time to remain in the 400-600 ms range. If the user desires a shorter plunge time for the same sample but still wants similar results, the glow discharge time can be approximately doubled. The opposite is true if needing to increase the plunge time range but wanting to obtain an overwicked grid.

Due to the mechanism for sample application by the chameleon to the grid surface – deposits in a stripe as the grid is moving vertically – a single sample stripe with an ice thickness gradient is obtained for each grid. We utilize this feature to locate the region of the stripe with the thinnest ice, ideally finding a couple of grid squares that have been overwicked sufficiently to no longer harbor thin films of sample across the holes, and then move to immediately adjacent grid squares with fewer empty holes and thin ice. We prioritize imaging these areas first since this will be the thinnest ice present in that grid, and then moving to adjacent areas until the ice thickness is no longer amenable to high-quality data collection and/or particle behaviors are sub-optimal. Notably, some particles will prefer slightly thicker ice or, depending on the shape of the molecule being imaged, different views of the molecule can be observed in different ice thicknesses due to orientation preferences. Knowing this, we routinely collect data from several adjacent grid squares (e.g., 2-5) as well as different areas within the holes of the same grid square (e.g., center versus closer to the hole edge) to obtain images of the particle of interest in a wider range of ice thicknesses (**Figure 9**). Using “on-the-fly” processing allows us to obtain real-time feedback on where high-quality data could be collected for each grid. Of course, not every EM grid prepared is worthy of collecting larger data sets necessary for high-resolution structure determination, so this scouting approach also affords the ability to “triage” grids much quicker than other blotting techniques that require much more scouting efforts to find areas of ideal ice thickness.

**Figure 9:**
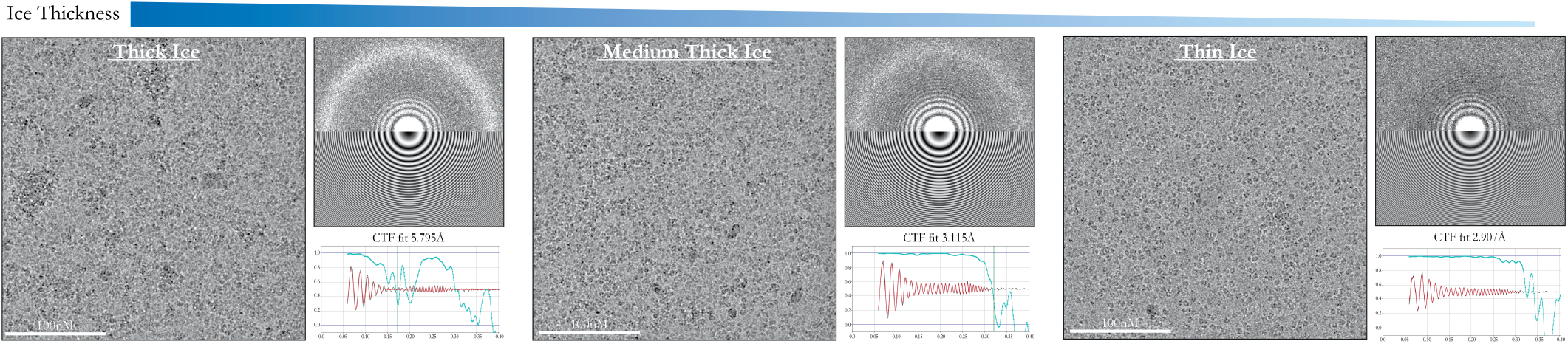
Contributions of ice thickness along the sample stripe to image quality. Micrographs of human hemoglobin resuspended in PBS frozen using a SPT Labtech chameleon using the settings described in this protocol. Three different areas along the sample ice stripe – thick, medium, and thin – are shown with corresponding contrast transfer function (CTF) estimation. Data were collected using EPU2 and processed using cryoSPARC Live version 4.2.1.

Ultimately, we recommend this scouting approach across a few grids prepared from the same freezing session but with different grid preparation parameters (e.g., glow discharge time, plunge time, etc.) to identify areas with the optimal particle density and ice thickness. If needed, we use these data to dial in the best parameters for a subsequent freezing session. We have also found it useful in a few cases to combine data collected across grids prepared using different glow discharge times and plunge times as critical views were observed in data collected from multiple grids, but not present in the data obtained from a single grid. Ultimately, the use of real-time grid wicking feedback with careful scouting in the electron microscope can produce high-quality data for structure determination.

## Acknowledgements

We thank the Herzik lab members, as well as UCSD’s CryoEM Facility director, Dr. Mariusz for critically thinking and providing feedback on this manuscript and the video content. We also would like to thank members of SPT Labtech, Tim Booth, Ouliana Panova, Paul Thaw, for guidance during manuscript preparation and operation of the chameleon. MAH is supported by NIH R35 GM138206 and as a Searle Scholar and a Cottrell Scholar. BDC is supported as a Goeddel Family Technology Sandbox Fellow. The chameleon used in this protocol was obtained via NIH 1S10OD032471 (MAH Co-I).

## References

1. Nakane, T.; Kotecha, A.; Sente, A.; McMullan, G.; Masiulis, S.; Brown, P.M.G.E.; Grigoras, I.T.; Malinauskaite, L.; Malinauskas, T.; Miehling, J.; et al. Single-particle cryo-EM at atomic resolution. Nature 2020, 587, 152–156. 10.1038/s41586-020-2829-0.

2. Li, X.; Zheng, S.Q.; Egami, K.; Agard, D.A.; Cheng, Y. Influence of electron dose rate on electron counting images recorded with the K2 camera. Journal of Structural Biology 2013, 184, 251–260. 10.1016/j.jsb.2013.08.005.

3. Brilot, A.F.; Chen, J.Z.; Cheng, A.; Pan, J.; Harrison, S.C.; Potter, C.S.; Carragher, B.; Henderson, R.; Grigorieff, N. Beam-induced motion of vitrified specimen on holey carbon film. Journal of Structural Biology 2012, 177, 630–637. 10.1016/j.jsb.2012.02.003.

4. McMullan, G.; Faruqi, A.; Henderson, R.; Guerrini, N.; Turchetta, R.; Jacobs, A.; van Hoften, G. Experimental observation of the improvement in MTF from backthinning a CMOS direct electron detector. Ultramicroscopy 2009, 109, 1144–1147. 10.1016/j.ultramic.2009.05.005.

5. Feathers, J.R.; Spoth, K.A.; Fromme, J.C. Experimental evaluation of super-resolution imaging and magnification choice in single-particle cryo-EM. Journal of Structural Biology: X 2021, 5, 100047. 10.1016/j.yjsbx.2021.100047.

6. Suloway, C.; Pulokas, J.; Fellmann, D.; Cheng, A.; Guerra, F.; Quispe, J.; Stagg, S.; Potter, C.S.; Carragher, B. Automated molecular microscopy: The new Leginon system. Journal of Structural Biology 2005, 151, 41–60. 10.1016/j.jsb.2005.03.010.

7. Cheng, A.; Negro, C.; Bruhn, J.F.; Rice, W.J.; Dallakyan, S.; Eng, E.T.; Waterman, D.G.; Potter, C.S.; Carragher, B. Leginon: New features and applications. Protein Science 2021, 30, 136–150. 10.1002/pro.3967.

8. Punjani, A.; Rubinstein, J.L.; Fleet, D.J.; Brubaker, M.A. cryoSPARC: algorithms for rapid unsupervised cryo-EM structure determination. Nat Methods 2017, 14, 290–296. 10.1038/nmeth.4169.

9. de la Rosa-Trevín, J.; Quintana, A.; del Cano, L.; Zaldívar, A.; Foche, I.; Gutiérrez, J.; Gómez-Blanco, J.; Burguet-Castell, J.; Cuenca-Alba, J.; Abrishami, V.; et al. Scipion: A software framework toward integration, reproducibility and validation in 3D electron microscopy. Journal of Structural Biology 2016, 195, 93–99. 10.1016/j.jsb.2016.04.010.

10. Zheng, S.Q.; Palovcak, E.; Armache, J.P.; Verba, K.A.; Cheng, Y.; Agard, D.A. MotionCor2: anisotropic correction of beam-induced motion for improved cryo-electron microscopy. Nat Methods 2017, 14, 331–332. 10.1038/nmeth.4193.

11. Dubochet, J.; McDowall, A.W.; Menge, B.; Schmid, E.N.; Lickfeld, K.G. Electron microscopy of frozen-hydrated bacteria. J Bacteriol 1983, 155, 381–390. 10.1128/jb.155.1.381-390.1983.

12. Dubochet, J.; McDowall, A. VITRIFICATION OF PURE WATER FOR ELECTRON MICROSCOPY. Journal of Microscopy 1981, 124, 3–4. 10.1111/j.1365-2818.1981.tb02483.x.

13. McDowall, A.W.; Chang, J.J.; Freeman, R.; Lepault, J.; Walter, C.A.; Dubochet, J. Electron microscopy of frozen hydrated sections of vitreous ice and vitrified biological samples. Journal of Microscopy 1983, 131, 1–9. 10.1111/j.1365-2818.1983.tb04225.x.

14. Nguyen, H.P.M.; McGuire, K.L.; Cook, B.D.; Herzik, Jr., M.A. Manual Blot-and-Plunge Freezing of Biological Specimens for Single-Particle Cryogenic Electron Microscopy. JoVE 2022, p. 62765. 10.3791/62765-v.

15. Dobro, M.J.; Melanson, L.A.; Jensen, G.J.; McDowall, A.W. Plunge Freezing for Electron Cryomicroscopy. In Methods in Enzymology; Elsevier, 2010; Vol. 481, pp. 63–82. 10.1016/S0076-6879(10)81003-1.

16. Resch, G.P.; Brandstetter, M.; Königsmaier, L.; Urban, E.; Pickl-Herk, A.M. Immersion Freezing of Suspended Particles and Cells for Cryo-Electron Microscopy. Cold Spring Harb Protoc 2011, 2011, pdb.prot5642. 10.1101/pdb.prot5642.

17. Resch, G.P.; Brandstetter, M.; Pickl-Herk, A.M.; Königsmaier, L.; Wonesch, V.I.; Urban, E. Immersion Freezing of Biological Specimens: Rationale, Principles, and Instrumentation. Cold Spring Harb Protoc 2011, 2011, pdb.top118. 10.1101/pdb.top118.

18. Lawson, C.L.; Baker, M.L.; Best, C.; Bi, C.; Dougherty, M.; Feng, P.; van Ginkel, G.; Devkota, B.; Lagerstedt, I.; Ludtke, S.J.; et al. EMDataBank.org: unified data resource for CryoEM. Nucleic Acids Research 2011, 39, D456–D464. 10.1093/nar/gkq880.

19. Noble, A.J.; Dandey, V.P.; Wei, H.; Brasch, J.; Chase, J.; Acharya, P.; Tan, Y.Z.; Zhang, Z.; Kim, L.Y.; Scapin, G.; et al. Routine single particle CryoEM sample and grid characterization by tomography. eLife 2018, 7, e34257. 10.7554/eLife.34257.

20. D’Imprima, E.; Floris, D.; Joppe, M.; Sánchez, R.; Grininger, M.; Kühlbrandt, W. Protein denaturation at the air-water interface and how to prevent it. eLife 2019, 8, e42747. 10.7554/eLife.42747.

21. Tan, Y.Z.; Baldwin, P.R.; Davis, J.H.; Williamson, J.R.; Potter, C.S.; Carragher, B.; Lyumkis, D. Addressing preferred specimen orientation in single-particle cryo-EM through tilting. Nat Methods 2017, 14, 793–796. 10.1038/nmeth.4347.

22. Rubinstein, J.L.; Guo, H.; Ripstein, Z.A.; Haydaroglu, A.; Au, A.; Yip, C.M.; Di Trani, J.M.; Benlekbir, S.; Kwok, T. Shake-itoff: a simple ultrasonic cryo-EM specimen-preparation device. Acta Crystallogr D Struct Biol 2019, 75, 1063–1070. 10.1107/S2059798319014372.

23. Feng, X.; Fu, Z.; Kaledhonkar, S.; Jia, Y.; Shah, B.; Jin, A.; Liu, Z.; Sun, M.; Chen, B.; Grassucci, R.A.; et al. A Fast and Effective Microfluidic Spraying-Plunging Method for High-Resolution Single-Particle Cryo-EM. Structure 2017, 25, 663–670.e3. 10.1016/j.str.2017.02.005.

24. Lu, Z.; Shaikh, T.R.; Barnard, D.; Meng, X.; Mohamed, H.; Yassin, A.; Mannella, C.A.; Agrawal, R.K.; Lu, T.M.; Wagenknecht, T. Monolithic microfluidic mixing–spraying devices for time-resolved cryo-electron microscopy. Journal of Structural Biology 2009, 168, 388–395. 10.1016/j.jsb.2009.08.004.

25. Dandey, V.P.; Wei, H.; Zhang, Z.; Tan, Y.Z.; Acharya, P.; Eng, E.T.; Rice, W.J.; Kahn, P.A.; Potter, C.S.; Carragher, B. Spotiton: New features and applications. Journal of Structural Biology 2018, 202, 161–169. 10.1016/j.jsb.2018.01.002.

26. Razinkov, I.; Dandey, V.P.; Wei, H.; Zhang, Z.; Melnekoff, D.; Rice, W.J.; Wigge, C.; Potter, C.S.; Carragher, B. A new method for vitrifying samples for cryoEM. Journal of Structural Biology 2016, 195, 190–198. 10.1016/j.jsb.2016.06.001.

27. Jain, T.; Sheehan, P.; Crum, J.; Carragher, B.; Potter, C.S. Spotiton: A prototype for an integrated inkjet dispense and vitrification system for cryo-TEM. Journal of Structural Biology 2012, 179, 68–75. 10.1016/j.jsb.2012.04.020.

28. Dandey, V.P.; Budell, W.C.; Wei, H.; Bobe, D.; Maruthi, K.; Kopylov, M.; Eng, E.T.; Kahn, P.A.; Hinshaw, J.E.; Kundu, N.; et al. Time-resolved cryo-EM using Spotiton. Nat Methods 2020, 17, 897–900. 10.1038/s41592-020-0925-6.

